# A convex optimization approach for identification of human tissue-specific interactomes

**DOI:** 10.1101/036830

**Authors:** Shahin Mohammadi, Ananth Grama

**Affiliations:** Department of Computer Sciences, Purdue University, West Lafayette, IN 47907, USA.

## Abstract

**Motivation:** Analysis of organism-specific interactomes has yielded novel insights into cellular function and coordination, understanding of pathology, and identification of markers and drug targets. Genes, however, can exhibit varying levels of cell-type specificity in their expression, and their coordinated expression manifests in tissue-specific function and pathology. Tissue-specific/ selective interaction mechanisms have significant applications in drug discovery, as they are more likely to reveal drug targets. Furthermore, tissue-specific transcription factors (tsTFs) are significantly implicated in human disease, including cancers. Finally, disease genes and protein complexes have the tendency to be differentially expressed in tissues in which defects cause pathology. These observations motivate the construction of refined tissue-specific interactomes from organism-specific interactomes.

**Results:** We present a novel technique for constructing human tissue-specific interactomes. Using a variety of validation tests (ESEA, GO Enrichment, Disease-Gene Subnetwork Compactness), we show that our proposed approach significantly outperforms state of the art techniques. Finally, using case studies of Alzheimer’s and Parkinson’s diseases, we show that tissue-specific interactomes derived from our study can be used to construct pathways implicated in pathology and demonstrate the use of these pathways in identifying novel targets.

**Availability:** http://www.cs.purdue.edu/homes/mohammas/projects/ActPro.html

## 1 Introduction

Proteins are basic workhorses of living cells. Their overall quantity is tightly regulated across different tissues and cell-types to manifest tissue-specific biology and pathobiology. These regulatory controls orchestrate cellular machinery at different levels of resolution, including, but not limited to, gene regulation (Mele *et al*., 2015; Göring, 2012), epigenetic modification (Chatterjee and Vinson, 2012; Mendizabal *et al*., 2014), alternative splicing (Buljan *et al*., 2012; Ellis *et al*., 2012), and post-translational modifications (Vaidyanathan and Wells, 2014; Ikegami *et al*., 2014). Transcriptional regulation is a key component of this hierarchical regulation, which has been widely used to study context-specific phenotypes. In the context of human tissues/ cell-types, genes can exhibit varying levels of specificity in their expression. They can be broadly classified as: (i) tissue-specific (unique to one cell-type); (ii) tissue-selective (shared among coherent groups of cell-types); and (iii) housekeeping (utilized in all cell-types). Tissue-specific/selective genes have significant applications in drug discovery, since they have been shown to be more likely drug targets (Dezso *et al*., 2008). Tissue-specific transcription factors (tsTFs) are significantly implicated in human diseases (Raj *et al*., 2014; Messina *et al*., 2004), including cancers (Vaquerizas *et al*., 2009). Finally, disease genes and protein complexes tend to be over-expressed in tissues in which defects cause pathology (Lage *et al*., 2008).

The majority of human proteins do not work in isolation but take part in pathways, complexes, and other functional modules. Tissue-specific proteins are known to follow a similar trend. Perturbations that impact interacting interfaces of proteins are significantly enriched among tissue-specific, disease-causing variants (Wang *et al*., 2012; Rolland *et al*., 2014; Sahni *et al*., 2015). This emphasizes the importance of constructing tissue-specific interactomes and their constitutive pathways for understanding mechanisms that differentiate cell-types and make them uniquely susceptible to tissue-specific disorders. Prior attempts at reconstructing human tissue-specific interactomes rely on a set of *“expressed genes”* in each tissue, and use this set as the baseline of transcriptional activity. The node removal (NR) method (Bossi and Lehner, 2009) constructs tissue-specific interactomes by identifying the induced subgraph of the expressed genes. Magger *et al*. (Magger *et al*., 2012) propose a method called *“Edge ReWeighting (ERW)”*, which extends the NR method to weighted graphs. This method penalizes an edge once, if one of its end-points is not expressed, and twice, if both end-points are missing from the expressed gene-set.

While these methods have been used to study tissue-specific interactions, their underlying construction relies only on the immediate end-points of each interaction to infer tissue-specificity. Furthermore, they threshold expression values, often using ad-hoc choices of thresholds to classify genes as either expressed or not. Finally, it is hard to integrate expression datasets from multiple platforms, or from multiple labs, into a single framework. These constraints are primarily dictated by limitations of high-throughput technologies for assaying gene expression. In these technologies, one can easily compare expression of the same gene across different samples to perform differential analysis; however, expression of different genes in the same sample are not directly comparable due to technical biases, differences in baseline expression, and GC content of genes. A recently proposed method, *Universal exPression Code (UPC)* (Piccolo *et al*., 2013), addresses many of these issues by removing platform-specific biases and converting raw expressions to a unified transcriptional activity score. These scores are properly normalized and can be compared across different genes and platforms.

Leveraging the UPC method, we propose a novel approach that uses the topological context of an interaction to infer its specificity score. Our approach formulates the inference problem as a suitably regularized convex optimization problem. The objective function of the optimization problem has two terms-the first term corresponds to a *diffusion kernel* that propagates activity of genes through interactions (network links). The second term is a *regularizer* that penalizes differences between *transcriptional* and *functional* activity scores. We use these functional activity scores to compute tissue-specificity for each edge in the global interactome, which we show, through a number of validation tests, are significantly better than prior methods. Our method is widely applicable and can be applied directly to single-channel, double-channel, and RNA-Seq expression datasets processed using UPC/SCAN. Furthermore, it can be easily adapted to cases where expression profiles are only available in preprocessed form.

The rest of the paper is organized as follows: In Section 2.1 we provide details of the datasets used in our study. Next, we introduce our method, called *Activity Propagation (ActPro)*, and provide a consistent notation to formalize previous methods. We evaluate the effectiveness of UPC transcriptional activity scores to predict tissue-specific genes in Section 3.1. Details of procedure for constructing tissue-specific networks and their parameter choices are discussed in Section 3.2. Section 3.3 provides qualitative assessment of our tissue-specific networks, whereas Sections 3.4-3.6 present validation studies for tissue-specific interactions using known pathway edges, co-annotation of proteins, and GWAS disease genes. Finally, in Section 3.7, we use the brain-specific interactome constructed using our method to identify novel disease-related pathways and use them to identify candidate targets for neurodegenerative disorders.

## 2 Materials and Methods

### 2.1 Datasets

We downloaded the RNASeq dataset version 4.0 (*dbGaP accession phs000424.v4.p1)* from the The Genotype-Tissue Expression (GTEx) project (Ardlie *et al*., 2015; Mele *et al*., 2015). This dataset contains 2,916 samples from 30 different tissues/cell types, the summary of which is presented in Figure 1. We processed each sample using the UPC method (Piccolo *et al*., 2013), a novel platform-independent normalization technique that corrects for platform-specific technical variations and estimates the probability of transcriptional activity for each gene in a given sample. The benefit of this method is that activation probability scores are highly consistent across different technologies, and more importantly, they are comparable across different genes in a given sample. For each gene, we recorded the transcript with the highest activation probability in the sample. Finally, we averaged replicate samples within each group to construct a unique transcription signature vector for each tissue/ cell type. The final dataset contains the expression value of 23,243 genes across 30 different tissues/ cell types.

**Fig. 1.**
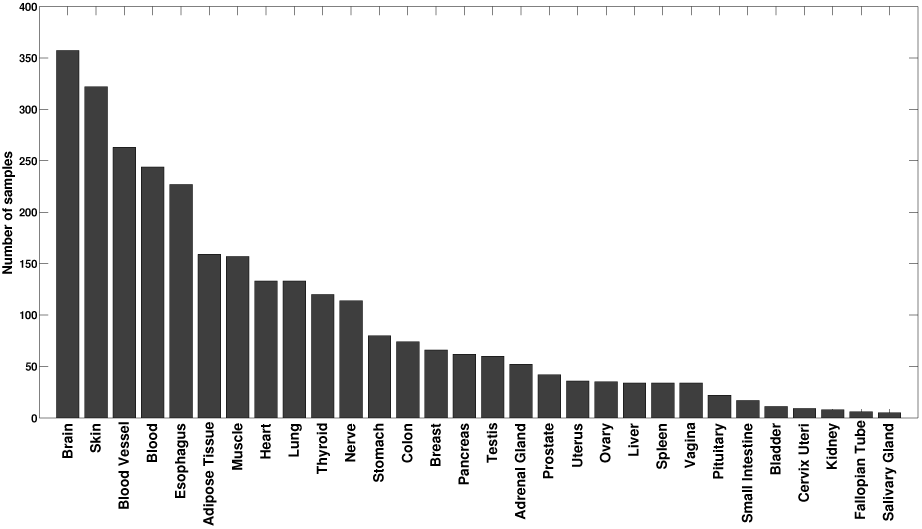
Summary of GTEx sample numbers per tissue.

In addition, we extracted human protein-protein interactions from the iRefIndex database (Razick *et al*., 2008), which consolidates protein interactions from different databases. Edges in this dataset are weighted using an MI (MINT-Inspired) score, which measures the confidence of each interaction based on three different evidence types, namely (i) the interaction types (binary/complex) and experimental method used for detection, (ii) the total number of unique PubMed publications reporting the interaction, and (iii) the cumulative evidence of interlogous interactions from other species. Finally, we map transcription data to the human interactome by converting all gene IDs to Entrez Gene IDs and only retaining genes that both have a corresponding node in the interactome and have been profiled by the GTEx project. This yields a global interactome with 147,444 edges, corresponding to protein-protein interactions, between 14,658 nodes, representing gene products.

### 2.2 Constructing human tissue-specific interactome

The global human interactome is a superset of all *possible* physical interactions that can take place in the cell. It does not provide any information as to which interactions actually occur in a given context. There are a variety of factors, including co-expression of genes corresponding to a pair of proteins, their co-localization, and post-translational modification, that mediate protein interactions at the right time and place. Quantifiable expression of both proteins involved in an interaction is one of the most important factors that determine the existence of an interaction. Different methods have been proposed in literature to utilize this source of information to construct human tissue-specific interactomes. Here, we briefly review existing methods, their drawbacks, and propose a new method, called *Activity Propagation (ActPro)*, which addresses noted shortcomings.

#### 2.2.1 Prior Methods

Let us denote the adjacency matrix of the global interatome by **A**, where element ***a***_*ij*_ is the weight (confidence) of the edge connecting vertices *v*_*i*_ and *v*_*j*_. Let ***z*** encode expression of genes in a tissue and *<u>**z**</u>* be the binarized version of ***z*** for a fixed threshold. Finally, let ***diag*** operator applied to a given vector be the diagonal matrix with the vector on the main diagonal. Our aim is to compute a matrix **Â**, which is the adjacency matrix of the tissue-specific interactome for a given expression profile. Using this notation, we can summarize prior methods for constructing tissue-specific interactomes as follows.

- **Node Removal (NR)** This method computes the induced subgraph of the “expressed” gene products (Bossi and Lehner, 2009).

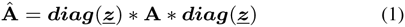
- **Edge Re-Weighting (ERW)** This method penalizes edges according to the expression state (active/inactive) of its end points (Magger *et al*., 2012). Given a penalty parameter 0 ≤ *rw* ≤ 1, ERW penalizes each edge by *rw* once, if only one of its end-points is active, and twice, if both incident vertices are inactive. Formally:

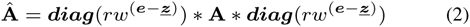

where ***e*** is the vector of all ones.

#### 2.2.2 Proposed Method

The main assumption of *ERW* and *NR* methods is that *transcriptional activity* of a gene is a reliable proxy for its *functional activity*. While this holds in a majority of cases, there are situations in which these scores differ significantly. First, the basis for *transcriptional activity* estimation is that genes with higher expression levels have higher chance of being functionally active in a given context. While this is generally true, there are genes that only need a low expression level to perform their function; i.e., their functionally active concentration is much lower than the rest of genes. Second, there is noise associated with measurement of gene expression, and converting measured expression values to UPC scores can over/ under-estimate *transcriptional activity*. Finally, we note that there are genes whose *down-regulation* corresponds to their functional activity (as opposed to the other way around).

Based on these observations, we propose a novel framework, called *Activity Propagation (ActPro)* to identify the most functionally active subnetwork of a given interactome. Our method incorporates global network topology to propagate activity scores, while simultaneously minimizing the number of changes to the gene activity scores. To this end, we first define a smoothed functional activity score defined by the following optimization problem:

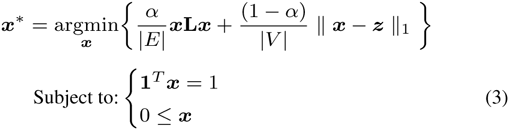

In this problem, **L** is the *Laplacian* matrix, defined as **A** – **Δ**, where element *δ*_*ii*_ of **Δ** is the weighted degree of *i*^*th*^ vertex in the global interactome. The Laplacian operator **L** acts on a given function defined over vertices of a graph, such as ***x***, and computes the smoothness of ***x*** over adjacent vertices. More specifically, we can expand the first term in 5 as Σ_*i,j*_ *w*_*i*,*j*_(*x*_*i*_ – *x*_*j*_)^2^, which is the accumulated difference of values between adjacent nodes scaled by the weight of the edge connecting them. This term defines a *diffusion kernel* that propagates activity of genes through network links. The second term is a *regularizer*, which penalizes changes by enforcing sparsity over the vector of differences between *transcriptional* and *functional* activities. This minimizes deviation from original transcriptome. It should be noted here that use of norm-1 is critical, since norm-2 regularization blends the transcriptional activity scores and significantly reduces their discriminating power. This negative aspect of norm-2 minimization is confirmed by our experiments. Finally, constraint **1**^*T*^***x*** = 1 is known as the fixed budget. It ensures that vector ***x*** is normalized and bounded. Parameter *α* determines the relative importance of regularization versus loss. We can equivalently define a penalization parameter 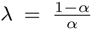 which is the standard notation in optimization framework. This problem is a classical convex optimization problem and we can solve it using efficient solvers to identify its global optimum.

After solving 5, we first scale ***x***^**\***^ by |*V*|. These scores are centered around 1, which allows us to perform minimal changes to the weight of interactions in the global interactome. Using these smoothed activity scores, we can re-weight the global human interactome as follows:

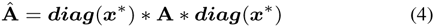

We can also derive an alternative formulation for *ActPro* which, instead of using *transcriptional activity* scores computed by UPC, uses expression values computed through more common methods such as RMA or MAS5.0 (Lim *et al*., 2007). We call this method *penalty propagation*, or *PenPro* for short. In this framework, computed expression values are not directly comparable and we need to threshold them to classify genes as either expressed or not. Using the same notation defined previously, we can define *functional activity* scores by solving the following problem:

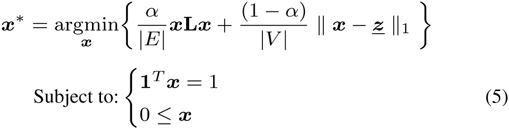

The only difference here is that, instead of *transcriptional activity* vector ***z***, we use the binarized expression vector ***<u>z</u>***. We observe similar performance for *ActPro* and *PenPro*, with *ActPro* being marginally superior in all cases, and thus we will only present results for *ActPro*.

### 2.3 Implementation details

All codes used in our experiments have been implemented in Matlab. To solve the convex problem in 5, we used CVX, a package for specifying and solving convex programs (Grant and Boyd, 2008). We used Mosek together with CVX, which is a high-performance solver for large-scale linear and quadratic programs (MOSEK-ApS, 2015).

## 3 Results and discussion

### 3.1 Transcriptional activity scores predict tissue-specificity of genes

To validate the quality of UPC normalized expression values, we first analyze the distribution of gene expressions across all tissues. Figure 2(a) shows the distribution of transcriptional activities, averaged over all samples. The overall distribution exhibits a bimodal characteristic that has a clear separation point that distinguishes expressed genes from others. We set a global threshold of 0.75 for identifying genes that are expressed in each tissue. These genes are used in evaluating NR and ERW methods. It should be noted that the distribution of UPC values vary across cell-types, as shown in Figure 2(b); however, the separation point is robust.

**Fig. 2.**
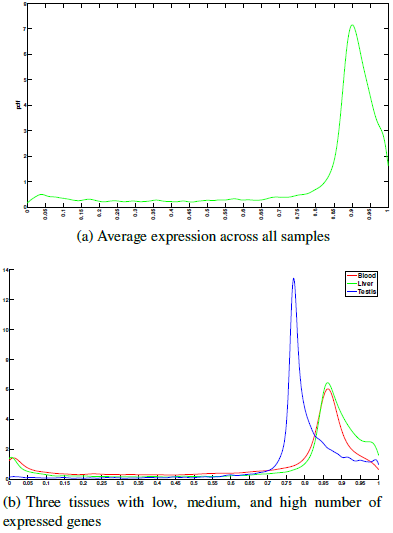
Distribution of UPC normalized gene expression values

Expression value of genes across tissues can be classified as *specific, selective*, or *housekeeping*. Housekeeping (HK) genes are ubiquitously expressed across all tissues to perform core cellular functions. On the other hand, tissue specific/selective genes are uniquely expressed in a given tissue context to perform tissue-specific functions. These genes typically reside in the periphery of the network, are enriched among signaling and cell surface receptors, and highly associated with the onset of tissue-specific disorders(Yeger-Lotem and Sharan, 2015). Figure 3(a) shows the total number of genes identified in each tissue as preferentially expressed (either specific or selective). Testis tissue exhibits the largest number of preferentially expressed genes (we refer to these as markers), with more that 1, 400 genes, while blood samples have the fewest markers with only ~ 250 marker genes. In order to assess whether the sets of preferentially expressed genes can predict tissue-specific functions, we performed GO enrichment analysis over different sets of tissue-specific markers using *GOsummaries* package in R/Bioconductor (Kolde, 2014). This package uses g:Profiler (Reimand *et al*., 2011) as backend for enrichment analysis and provides a simple visualization of the results as a word cloud. The coverage of available annotations for different tissues is not uniform; that is, some tissues are better annotated for specific terms than the others. We chose six well-annotated tissues with high, mid, and low number of identified markers for further study. We limited terms to the ones with at least 20 and at most 500 genes to avoid overly generic/specific terms. Finally, we used a strong hierarchical filtering to remove duplicate GO terms and thresholded terms at *p*-value of 0.05. Figure 3(b) shows the enrichment word-cloud for each tissue. It can be seen that all terms identified here are highly tissue-specific and representative of main functions for each tissue, which supports the validity of computed transcriptional activity scores from UPC.

**Fig. 3.**
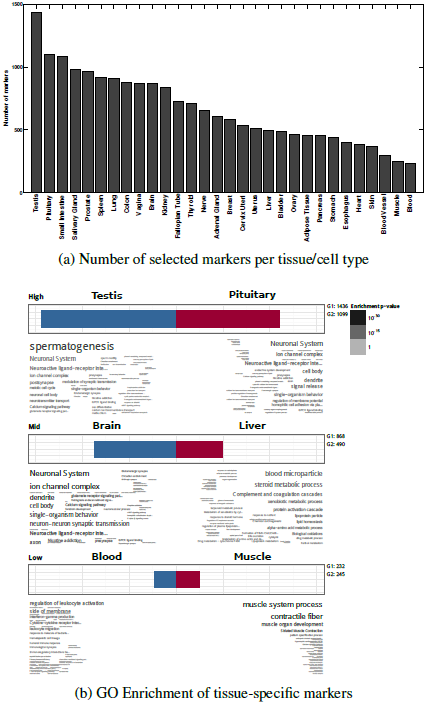
Evaluation of tissue-specific markers using a threshold value of 0.75 to define expressed genes.

### 3.2 Constructing tissue-specific interactomes

Node Removal (NR) and Edge ReWeighting (ERW) methods need a predefined set of expressed genes in each tissue to construct tissue-specific interactomes (or a given lower bound to threshold expression values). We use the set of all genes with transcriptional activity greater than or equal to 0.75 as the set of expressed genes for these methods. We chose this threshold based on the averaged distribution of gene expressions, as well as further manual curation of genes at different thresholds.

Node Removal (NR) method is known to disintegrate the network with stringent expression values(Magger *et al*., 2012). To evaluate the performance of NR over different expression thresholds and assess its sensitivity to the choice of threshold, we computed the size of largest connected components, while varying the value of expression threshold. Figure 4 shows stable behavior up to threshold value of 0.75, after which the size of largest component exhibit a rapid shift and the network starts to disintegrate. This suggests that the expression value of 0.75 is also the optimal topological choice for NR method.

**Fig. 4.**
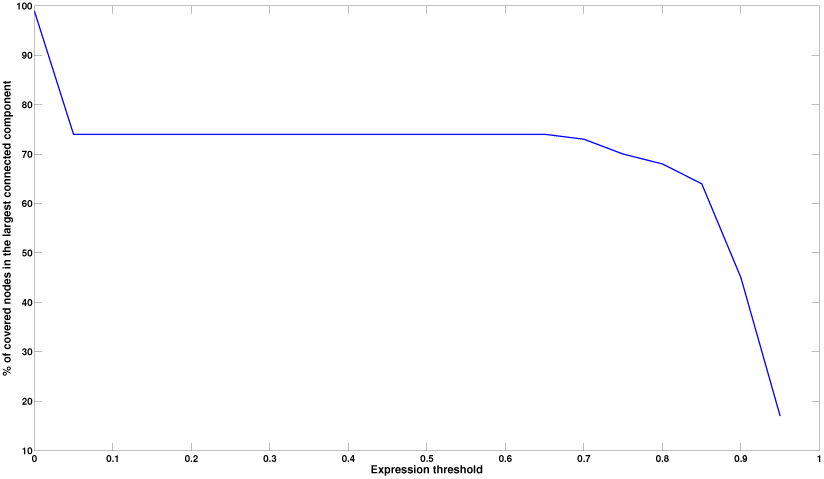
Size of the largest connected component in node removal (NR) method as a function of expression threshold. A rapid disaggregation phase can be spotted around 0.75

For the ActPro algorithm, we evaluated the results over three different values of *α* in set {0.15, 0.5, 0.85} and reported the result for each case.

### 3.3 Qualitative characterization of tissue-specific interactomes

A key feature of tissue specific networks is their ability to discriminate positive edges that manifest in each tissue from the entire set of potential interactions in the global interactome. In case of Node Removal (NR) and Edge ReWeighting (ERW) methods, it is easy to distinguish positive and negative edges: every edge for which at least one of the endpoints is not expressed can be classified as a negative edge. The latter method updates edge weights, to account for expression of their end-points, whereas the former method sets a hard threshold to either include an edge or not. In the case of *ActPro*, we first notice that the distribution of edge weights is very different between ActPro and previous methods. Whereas NR and ERW methods never increase the weight of an edge, in ActPro edge weights can increase or decrease. This behavior, however, is biased towards the positive end. To decompose each network into its HK, positive, and negative subspaces, we use the following strategy: for each tissue-specific network constructed by a given method, we first compute the relative weight change between the global interactome and the tissue-specific network. We then normalize these changes using Z-score normalization and define positive and negative subspaces according to the sign of normalized relative changes. We further define and separate HK edges as the subset of positive edges that are positive in at least half of the tissues. Figure 5 summarizes the average statistics for constructed networks using different methods. As a general observation, *ActPro* classifies fewer interactions as housekeeping and provides more specific positive and negative edges. Furthermore, as we increase the *α* parameter, representing the diffusion depth, we observe that these edges are more evenly distributed across vertices. To give a concrete example, we constructed the brain-specific network using *ERW* and *ActPro* methods. Figure 6 illustrates the final statistics of the constructed networks. Consistent with the average statistics, we observe much smaller positive/negative nodes/edges in *ERW*.

**Fig. 5.**
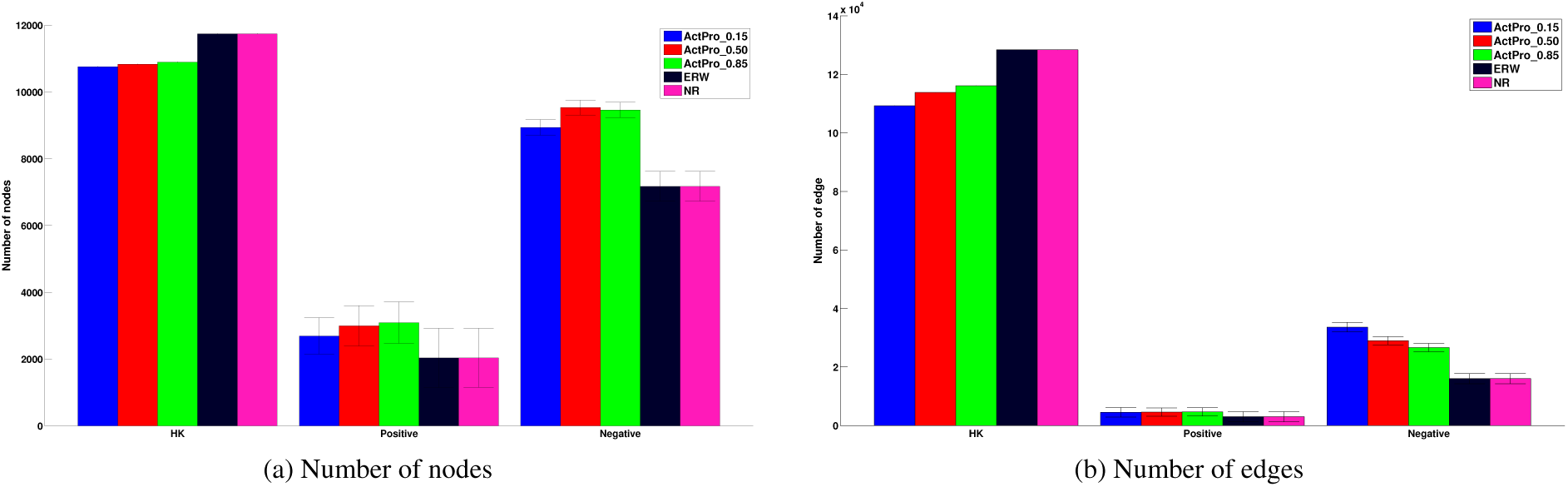
Qualitative characteristics of tissue-specific interactomes constructed using different methods. Housekeeping (HK) subset is shared across all tissues constructed by a given method, whereas positive subset is unique to each tissue. Negative subset contains interactions that are not utilized in a given tissue.

**Fig. 6.**
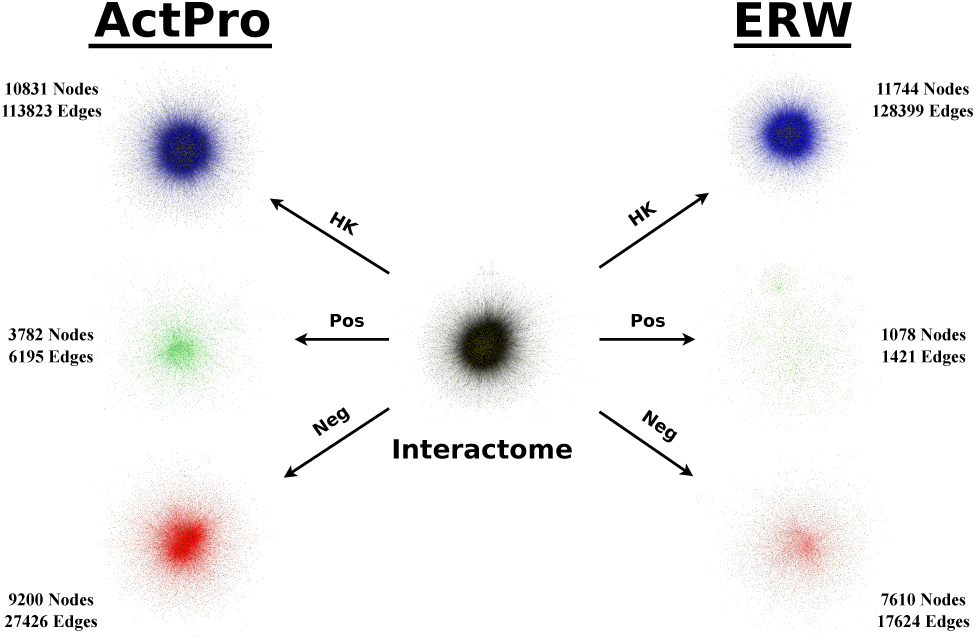
Decomposition of global interactome into brain-specific network using ERW and ActPro (*α* = 0.5) methods

### 3.4 Tissue-specific interactome predicts context-sensitive interactions in known functional pathways

To evaluate the power of tissue-specific interactions in capturing context-sensitive physical interactions in known pathways, we first use Edge Set Enrichment Analysis (ESEA) to rank pathway edges according to their gain/loss of mutual information in each tissue context (Han *et al*., 2015). ESEA aggregates pathways from seven different sources (KEGG; Reactome; Biocarta, NCI/Nature Pathway Interaction Database; SPIKE; HumanCyc; and Panther) and represents them as a graph with edges corresponding to biological relationships, resulting in over 2,300 pathways spanning 130,926 aggregated edges. It then uses an information-theoretic measure to quantify dependencies between genes based on gene expression data and ranks edges, accordingly. Formally, for each pathway edge, ESEA computes the differential correlation score (EdgeScore) as follows:

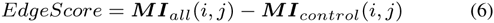

where ***MI***_*all*_ is the mutual information of the gene expression profiles for genes *i* and *j* across all cell-types. Here, ***MI***_*control*_ measures the mutual information only in the given tissue context. Each edge can be classified as either a gain of correlation (GoC), loss of correlation (LoC), or no change (NC) depending on the value of *EdgeScore*. We use GoC edges, that is, a pair of genes with positive gain of mutual information in the tissue context, as true positive edges in each tissue. Similarly, we use all positive edges in all tissues but the tissue of interest as true negatives.

To assess agreement between ESEA scores over known pathway edges and computed tissue-specific interactions, we rank all edges according to the difference of their weights in the human tissue-specific interactome compared to the global interactome and evaluate the enrichment of true positive pathway edges among top-ranked edges.We compute the receiver operating characteristic (ROC) curve for each tissue and average the area under the curve (AUC) gain, compared to random baseline, over all tissues. Figure 7 presents the relative performance of each method. All three configurations of the *ActPro* algorithm are ranked at the top of the list – demonstrating the superior performance of our proposed method.

To further investigate tissue-specific details for the top-ranked method, *ActPro* with *α* = 0.15, we sorted AUC gain for each tissue, shown in Figure 8. This plot exhibits high level of heterogeneity, and surprisingly, four of the tissues had worse than random performance. This was consistent across all of the methods. To further understand this, we investigated the ranked list of edges and identified a high enrichment of edges with LoC among top-ranked edges. We performed enrichment analysis over these negative edges and identified significant tissue-specific functions among them, which suggests that the poor observed performance for these tissues is attributed to their misclassification as negative edges.

**Fig. 7.**
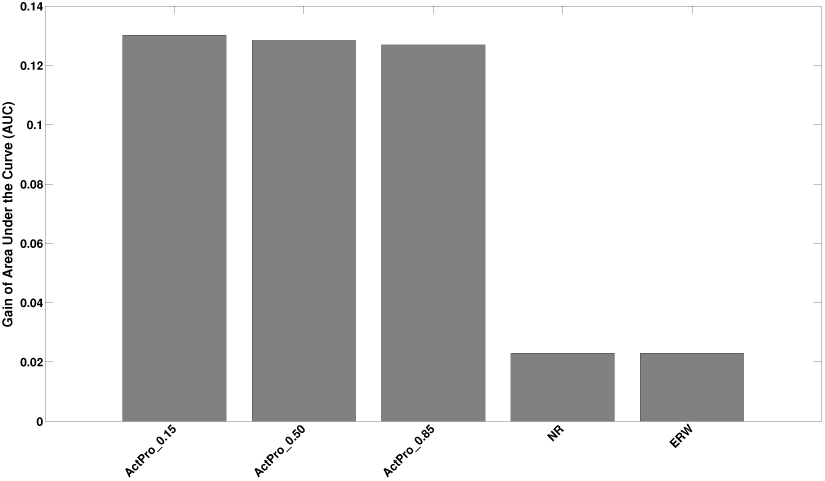
Gain of Area Under the Curve (AUC) of known context-specific pathway edges among tissue-specific interactions

**Fig. 8.**
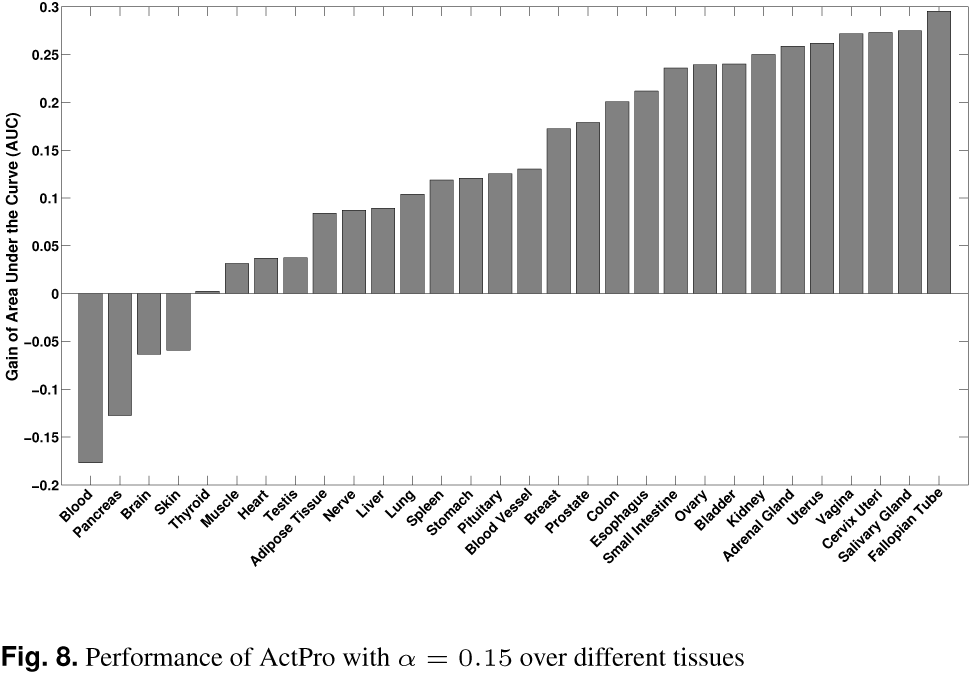
Performance of ActPro with *α* = 0.15 over different tissues

At the other end of the spectrum, *Fallopian Tube, Vagina*, and *Cervix Uteri* had consistently high AUC gain across different methods. Figure 9 shows the ROC curve for these tissues.

**Fig. 9.**
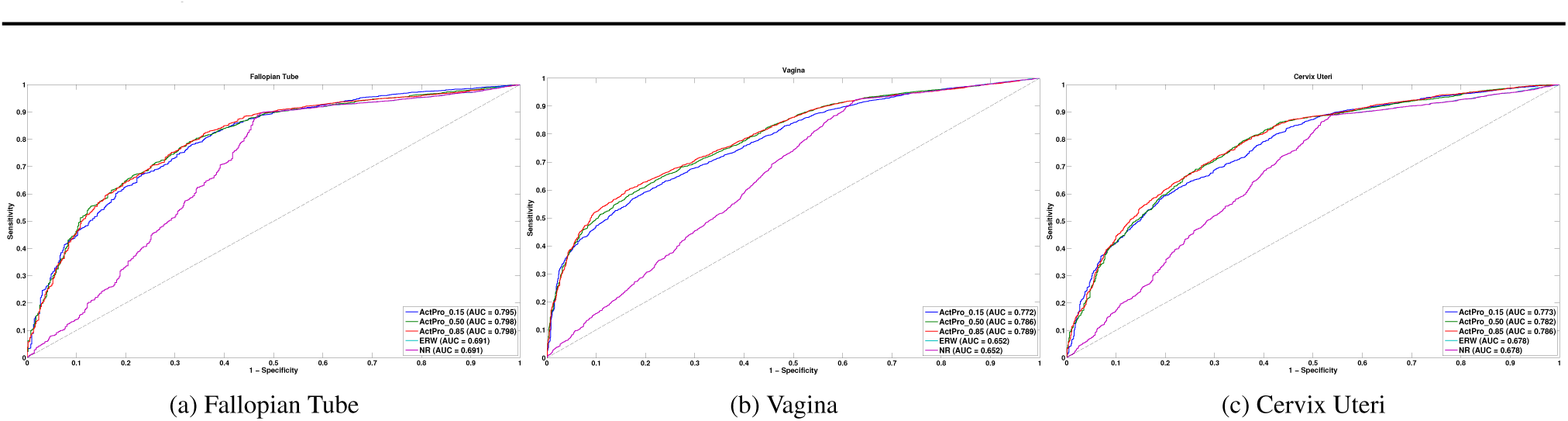
Tissues with the highest gain of AUC for predicting tissue-specific pathway edges

### 3.5 Tissue-specific interactions are enriched among proteins with shared tissue-specific annotations

We hypothesize that tissue-specific edges are enriched with proteins that participate in similar tissue-specific functions. To evaluate our hypothesis, we collected a set of manually curated tissue-specific Gene Ontology (GO) annotations from a recent study (Greene *et al*., 2015). We mapped tissues to GTEx tissues and identified tissue-specific GO annotations for genes in each tissue-specific interactome. We excluded tissues with less that 100 edges with known annotations. This resulted in 10 tissues, *Adipose Tissue, Blood Vessel, Blood, Brain, Breast, Heart, Kidney, Lung*, and *Muscle*, for which we had enough annotations. We use the same strategy employed in previous section to identify the mean gain of AUC for each method, which is illustrated in Figure 10. It should be noted that the gain of AUC is much smaller here than the case with ESEA edges, which can be attributed to the sparsity of tissue-specific GO annotations. Unlike ESEA, *ActPro* with *α* = 0.5 outperforms the case with *α* = 0.15.

**Fig. 10.**
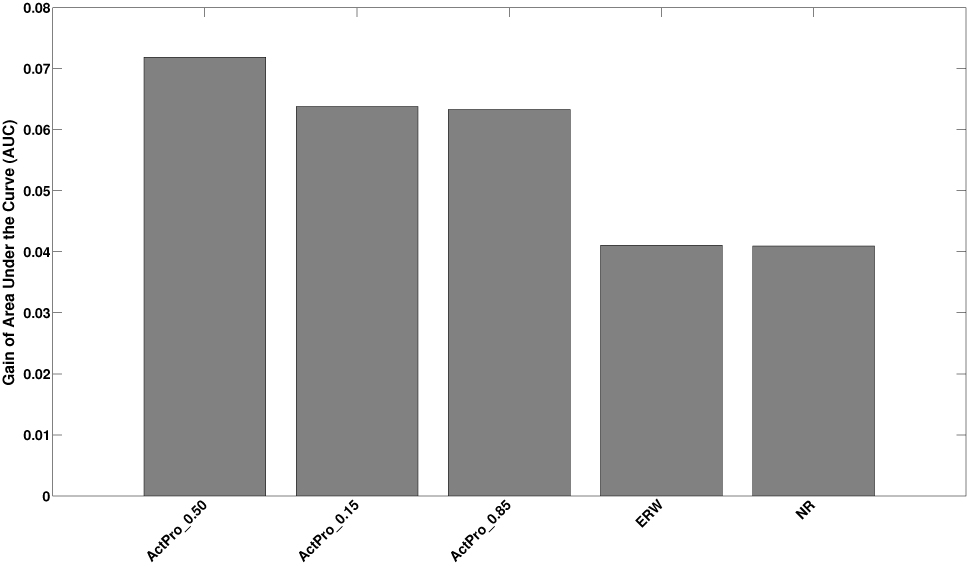
Mean gain of Area under the curve (AUC) for predicting proteins co-annotated with tissue-specific functions

Among the ten tissues, *Adipose* and *Muscle* tissues performed marginally better than the others with AUC of 0.59 and 0.58, respectively. On the other hand, *Lung* tissue had the worst performance with lower than random AUC of 0.47.

### 3.6 Tissue-specific interactions densely connect genes corresponding to tissue-specific disorders

Disease genes are densely connected to each other in the interactome, which provides the basis for a number of methods for network-based disease gene prioritization (Köhler *et al*., 2008). Tissue-specific interactomes have been shown to have higher accuracy in predicting disease-related genes using the random-walk method (Magger *et al*., 2012). More recently, Cornish *et al*. (Cornish *et al*., 2015) used the concept of “geneset compactness”, and showed that the average distance among nodes corresponding to a given disorder is significantly smaller in tissue-specific networks, compared to an ensemble of random graphs.

Here, we adopt this concept to measure how closely tissue-specific genes related to human disorders are positioned in networks constructed using different methods. First, we use a symmetric diffusion process instead of Random-Walk with Restart (RWR), which is a better measure of distance. Second, we use an alternative random model in which we hypothesize that genes corresponding to tissue-specific disorders are strongly connected to each other, compared to random genesets of the same size.

To validate our hypothesis, we gather genes corresponding to tissue-specific disorders from a recent study (Himmelstein and Baranzini, 2015). These genes are extracted from the GWAS Catalog by mapping known associations to disease-specific loci. Among a total of 99 disorders, we focused on the gold standard set of 29 diseases with at least 10 high-quality primary targets. We successfully mapped 27 of these diseases to GTEx tissues, which are used for the rest of our study. Consistent with previous studies (Magger *et al*., 2012), we observed a small subset of disease genes not to be expressed in the tissue in which they cause pathology. Among all disease genes, we only retained genes that are connected in the global interactome and are expressed above 0.1 UPC score.

For a given tissue-specific interactome represented by its adjacency matrix, ***A***_*T*_, we define a stochastic matrix 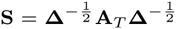, where **Δ** is the diagonal matrix, with entries *δ*_*ii*_ being the degree of node *i* in the human tissue-specific interactome. Using this matrix, we can compute degree-weighted random-walk scores among gene pairs as:

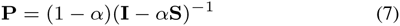

We define the random-walk distance as *d*_*ij*_ = –*log*_10_(*p*_*ij*_), after replacing zero elements of **P** with *ϵ* = 2^−52^. Given a disease geneset Γ, we measure its compactness as the normalized average of distances for all pairs of nodes in the geneset:

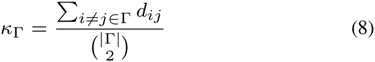

Finally, we sampled without replacement, 100K vertex samples of size |Γ| from the tissue-specific interactome and computed the compactness for each of the samples, individually. We defined an empirical *p*-value as the fraction of random instances with higher compactness (lower *κ*) compared to Γ. We removed disorders for which none of the methods yield significant *p*-value given a threshold of 0.05. The final dataset consists of 15 diseases with significantly compact interactions. To combine the *p*-values for different disorders, we use the Edgington method (Edgington, 1972). This method gathers a statistic 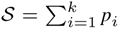 for a set of *k* given *p-values*, and computes the meta *p*-value by assigning significance to *S* as:

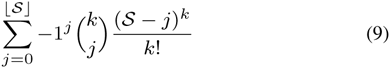

The list of all individual and combined p-values is shown in Table 1. In these experiments, *ActPro* (*α* = 0.85) had the most significant results, closely followed by *ActPro* (*α* = 0.5). This suggests that propagating information using diffusion kernel in ActPro enhances its prediction power for tissue-specific pathologies. Furthermore, there are four diseases for which the global interactome had more significant predictions compared to tissue-specific networks, among which *primary biliary cirrhosis* and *psoriasis* had the highest difference. This difference may be attributed to misclassification of disease/ tissue in Himmelstein and Baranzini (2015), or existence of cross tissue mechanisms of action for the disease.

### 3.7 Tissue-specific interactome identifies novel disease-related pathways – case study in neurodegenerative disorders

We now investigate whether tissue-specific interactomes can help in predicting novel pathways that are involved in the progression of neurodegenerative disorders. We perform a case study of *Alzheimer’s* and *Parkinson's* diseases, both of which were among disorders with high compactness in brain tissue. We use Prize-Collecting Steiner Tree (PCST) algorithm to identify the underlying pathway among disease-genes identified by GWAS studies. Formally, PCST problem can be formulated as:

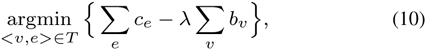

where *T* is an induced tree of the given graph, *v* and *e* are the set of vertices and edges in *T*, respectively, *c*_*e*_ is the cost of choosing edge *e*, and *b*_*v*_ is the reward/prize of collecting node *v*. Similar methods have been proposed previously to connect upstream signaling elements to downstream transcriptional effector genes (Tuncbag *et al*., 2012, 2013).

To identify disease-related pathways, we first prune non-specific interactions in the network by removing vertices that have more than 500 interactions. We transform edge confidence values (conductances) to edge penalties (resistances) by inverting each edge weight. Node prizes are defined as the ratio of their incident edges that fall within disease-related genes to the total degree of a node. We assigned a node prize of 1,000 to disease genes to ensure that they are selected as terminal nodes. Finally, we use a recent message passing algorithm (Bailly-Bechet *et al*., 2011) to identify PCST rooted at each disease-related gene and choose the best tree as the backbone of the disease-related pathway. Over each node, we use a maximum depth of 4 and λ = 1 as parameters to the message passing algorithm. Figure 11 shows final tissue-specific pathways for Alzheimer’s and Parkinson’s diseases.

Alzheimer’s disease (AD) network contains two distinct subnetworks, one centered around *CLTC* and the other centered around *ABL1. PICALM, CLU, APOE*, and *SORL1* are all known genes involved in AD, which are also involved *negative regulation of amyloid precursor protein catabolic process*. All four of these genes converge on *CLCT* gene, but through different paths. *PICALM* gene is known to play a central role in clathrin-related endocytosis. This protein directly binds to CLTC and recruits clathrin and adaptor protein 2 (AP-2) to the plasma membrane (Carter, 2011). On the other hand, *CLU, APOE*, and *SORL1* are linked to the *CLTC* through novel linker genes *XRCC6, MAPT/BIN1* and *GG2A/HGS*, respectively. Gamma-adaptin gene, GGA2, binds to clathrins and regulates protein traffic between the Golgi network and the lysosome. This network is postulated to be an important player in AD (Carter, 2011). *HGS* gene is a risk factor age-related macular degeneration (AMD) and has been hypothesized to be a shared factor for AD (Logue *et al*., 2014).

**Table 1.** Compactness of tissue specific disease genes in their tissue-specific interactome

|  | global | ActPro_0.15 | ActPro_0.50 | ActPro_0.85 | ERW | NR |
| --- | --- | --- | --- | --- | --- | --- |
| Alzheimer's disease | **4.12E-3** | 6.96E-3 | 5.98E-3 | 5.44E-3 | 5.32E-3 | 9.60E-2 |
| breast carcinoma | 1.83E-3 | 1.11E-3 | 8.40E-4 | **8.30E-4** | 4.09E-3 | 8.15E-2 |
| chronic lymphocytic leukemia | 8.20E-4 | 7.40E-4 | **4.80E-4** | 5.10E-4 | 8.50E-4 | 2.94E-2 |
| coronary artery disease | 3.95E-1 | 1.58E-1 | 1.09E-1 | 1.03E-1 | 1.33E-1 | **1.93E-2** |
| Crohn's disease | 2.56E-2 | 1.93E-2 | 1.50E-2 | **1.44E-2** | 8.54E-2 | 4.14E-1 |
| metabolic syndrome X | 1.11E-2 | 1.09E-2 | **1.07E-2** | 1.12E-2 | 1.02E-1 | 7.39E-1 |
| Parkinson's disease | 1.59E-2 | 1.25E-2 | 9.89E-3 | **9.50E-3** | 1.34E-2 | 9.62E-2 |
| primary biliary cirrhosis | **7.20E-4** | 1.32E-3 | 3.16E-3 | 3.40E-3 | 2.80E-2 | 6.86E-1 |
| psoriasis | **2.10E-4** | 1.10E-3 | 1.16E-3 | 9.50E-4 | 4.67E-3 | 3.24E-1 |
| rheumatoid arthritis | 1.70E-2 | **9.28E-3** | 1.06E-2 | 1.10E-2 | 6.39E-2 | 3.61E-1 |
| systemic lupus erythematosus | 4.98E-2 | 1.19E-2 | 7.56E-3 | 7.22E-3 | 2.55E-3 | **1.60E-4** |
| type 1 diabetes mellitus | 2.64E-2 | 3.01E-2 | **2.38E-2** | 2.40E-2 | 2.64E-1 | 9.39E-1 |
| type 2 diabetes mellitus | 1.57E-3 | 2.90E-4 | 2.40E-4 | **1.80E-4** | 5.60E-4 | 7.90E-3 |
| vitiligo | **1.17E-3** | 2.13E-3 | 3.04E-3 | 3.54E-3 | 1.84E-2 | 5.69E-1 |
| schizophrenia | 3.47E-1 | 2.13E-1 | 1.93E-1 | 1.84E-1 | 1.40E-1 | **4.10E-2** |
| combined | 1.53E-13 | 1.24E-17 | 6.62E-19 | **3.70E-19** | 9.03E-14 | 2.43E-03 |

**Fig. 11.**
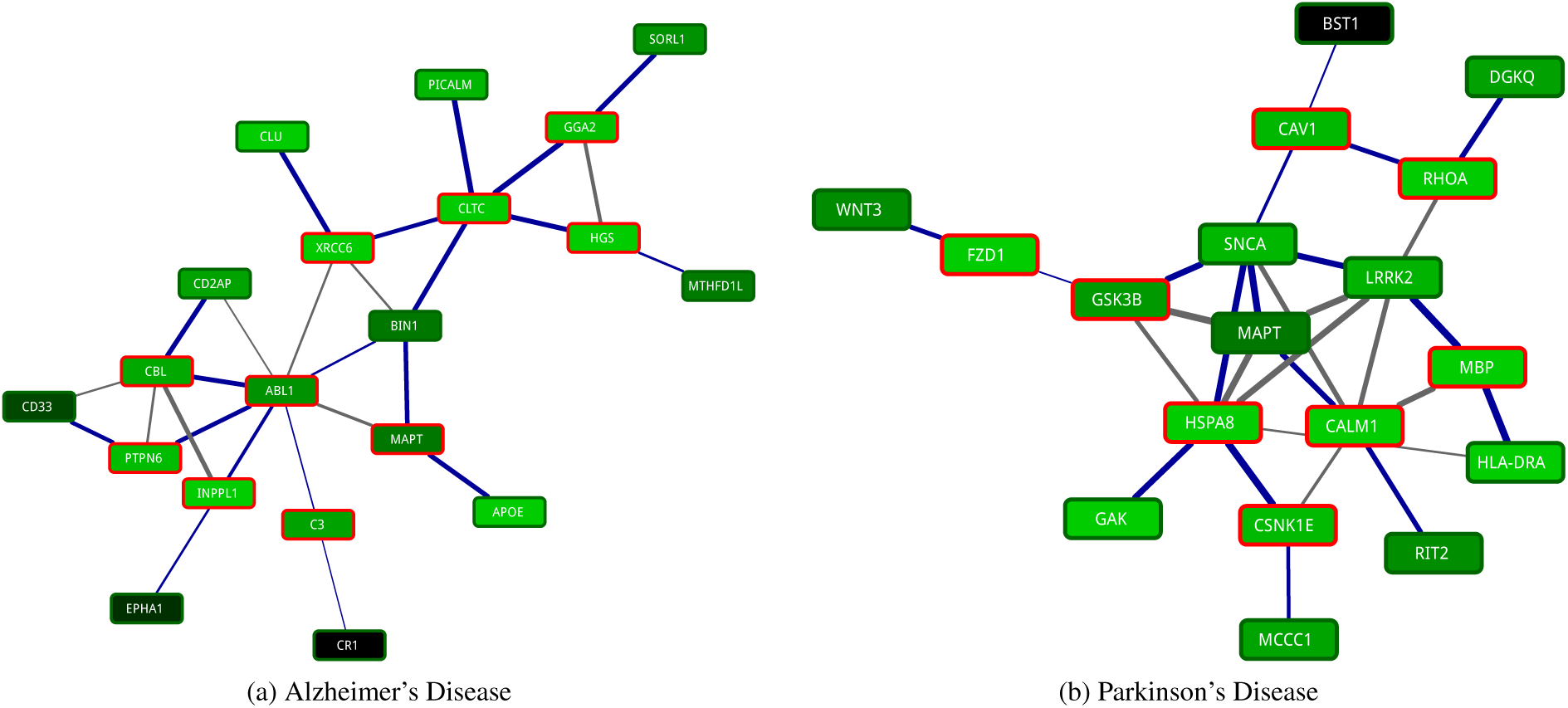
Tissue-specific pathways in human neurodegenerative disorders. Nodes are colored according to their tissue-specific expressions, with novel identified genes marked in red, accordingly. The thickness of edges represent their confidence with tree edges marked as blue.

Interestingly, *MAPT*, a novel marker identified in this study, is a risk factor for Parkinson’s disease and very recently shown to also be linked to AD (Desikan *et al*., 2015). A second component in AD network is centered around *ABL1* gene, which, together with *CBL, INPPL1, CD2AP*, and *MAPT* share the *SH3 domain binding* function. *INPPL1* gene, a metabolic syndrome risk factor, has been hypothesized to link AD with the recently posed term *“type 3 diabetes”* (Accardi *et al*., 2012). Finally, we note that *MAPT* gene is one of the central genes that links these two main components, the role of which warrants further investigation.

Parkinson’s disease (PD) network, on the other hand, contains one densely connected core centered around *MAPT* gene. There are two main branches converging on *MAPT*. On the left, *WNT3, FZD1*, and *GSK3B* constitute upstream elements of the WNT signaling pathway, which is known to play an important role in PD neurodegeneration (Berwick and Harvey, 2012). *GSK3* gene product is postulated to directly interact with *MAPT* (*τ*) and *LRKK2*, while implicitly regulating *SNCA* (*α*-Syn) in a *β*-cat dependent manner. However, we observed direct interaction between *GSK3B* and **SNCA**, and parallel pathways connecting it to *LRRK2* via

*SNCA* and *MAPT*. Both *SNCA* and *MAPT* also take part in the right branch, together with *CAV1* and *RHOA*, which is enriched in *reactive oxygen species metabolic process*. Accumulation of ROS contributes to mitochondrial dysfunction and protein misfolding, both of which are linked to progression of PD. *RIT2* enzyme is identified independently and confirmed as PD susceptibility factor (Pankratz *et al*., 2012). Pankratz *et al*. also suggested *CALM1* as the bridge linking *RIT2* with *MAPT* and *SNCA*, which confirms our findings. Cyclin G associated kinase (*GAK*) is a known risk factor for PD. We identified *HSPA8* as a key link between *GAK*, WNT signaling pathway, and *CSNK1E* with central PD genes, *MAPT, SNCA*, and *LRRK2*. *HSPA8* gene has been proposed as a biomarker for diagnosis of PD (Lauterbach, 2013). Finally, myelin basic protein (*MBP*) interacts closely with *CALM1* and *LRRK2*. This gene has been previously shown to be differentially expressed in PD and proposed as potential biomarker for PD (Kim *et al*., 2006).

In summary, we show that the brain-specific interactome derived from our method helps in uncovering tissue-specific pathways that are involved in neurodegenerative diseases. Similar analysis of other human tissues can potentially contribute to identification of new therapeutic targets for other human disorders.

## 4 Conclusion

In this paper, we present a novel method for computing tissue-specific interactomes from organism-specific interactomes and expression profiles of genes in various tissues. Our method casts the problem as a convex optimization problem that diffuses functional activity of genes over the organism-specific interactome, while simultaneously minimizing perturbation of transcriptional activity. We show, using a number of validation studies, that the tissue-specific interactomes computed by our method, are superior to those computed using existing methods. Finally, we show, using a case study of brain-specific interactome for Alzheimer’s and Parkinson’s diseases, that our method is capable of constructing highly resolved disease-specific pathways, providing potential targets for novel drugs.

